# The wall-less bacterium *Spiroplasma poulsonii* builds a polymeric cytoskeleton composed of interacting MreB isoforms

**DOI:** 10.1101/2021.06.08.447548

**Authors:** Florent Masson, Xavier Pierrat, Bruno Lemaitre, Alexandre Persat

## Abstract

A rigid cell wall defines the morphology of most bacteria. MreB, a bacterial homologue of actin, plays a major role in coordinating cell wall biogenesis and defining a cell’s shape. In contrast with most bacteria, the Mollicutes family is devoid of cell wall. As a consequence, many Mollicutes have undefined morphologies. *Spiroplasma* species are an exception as they robustly grow with a characteristic helical shape, but how they maintain their morphology remains unclear. Paradoxal to their lack of cell wall, the genome of *Spiroplasma* contains five homologues of MreB (SpMreBs). Since MreB is a homolog of actin and that short MreB filaments participate in its function, we hypothesize that SpMreBs form a polymeric cytoskeleton. Here, we investigate the function of SpMreB in forming a polymeric cytoskeleton by focusing on the Drosophila endosymbiont *Spiroplasma poulsonii*. We found that *in vivo, Spiroplasma* maintain a high concentration of all five MreB isoforms. By leveraging a heterologous expression system that bypasses the poor genetic tractability of *Spiroplasma*, we found that strong intracellular levels of SpMreb systematically produced polymeric filaments of various morphologies. Using co-immunoprecipitation and co-expression of fluorescent fusions, we characterized an interaction network between isoforms that regulate the filaments formation. Our results point to a sub-functionalization of each isoform which, when all combined *in vivo*, form a complex inner polymeric network that shapes the cell in a wall-independent manner. Our work therefore supports the hypothesis where MreB mechanically supports the cell membrane, thus forming a cytoskeleton.

**Significance statement:** Bacteria shape is determined by their cell wall. The actin homologue MreB essentially determines shape by organizing cell wall synthesis at the subcellular level. Despite their lack of cell wall, *Spiroplasma* robustly grow into long helical bacteria. Surprisingly, its genome retains five copies of *mreB* while it lost genes encoding canonical MreB interactors. We sought to delineate the exact function of *Spiroplasma* MreBs (SpMreBs). We leveraged *in vivo* data along with functional studies to systematically investigate MreB polymerization behavior. We uncovered that SpMreBs build into filaments, which structure it determined by a complex interaction network between isoforms. Our results support the hypothesis that MreB can mechanically support the membrane of *Spiroplasma*, hence acting as a load-bearing cytoskeletal protein.

## Introduction

Bacteria grow into an astonishing variety of shapes including spheres, straight and curved rods, disks, trapezoids, helices and even stars (1). The stability of these morphologies within each species suggests that they confer important fitness advantages in the natural ecological niches of these microbes (2). Bacteria typically maintain their morphologies by virtue of their rigid cell wall (3). Local constraining of cell wall peptidoglycan patterning by short MreB polymers provides a cell with its shapes (4). As a consequence, degrading the bacterial cell wall is sufficient to relax its shape into a spherical membrane (5, 6). In contrast, most mamalian cells are plastic and actively control their shapes by virtue of a dynamic actin cytoskeleton (7).

All but one bacterial family (Mollicutes) synthesize a peptidoglycan cell wall. The cell envelope of Mollicutes is essentially a lipid bilayer. Mollicutes underwent a genome reduction, including loss of cell wall synthesis genes, by regressive evolution upon their adaptation to a strict host-associated lifestyle (8, 9). As one could expect from their lack of cell wall, the class encompasses genera such as *Mycoplasma* and *Phytoplasma* with no distinctive cell shape, adapting their morphology to the constraints of their close environment (10). *Spiroplasma* makes exception as this genus is uniformly composed of long, helical bacterial species (11) (Figure 1A and Supplemental movie 1). Furthermore, *Spiroplasma* cells actively deform their body to propel themselves in high viscosity fluid environments (12, 13). How does *Spiroplasma* maintain and actively controls its morphology? No external rigid structure has been identified in these bacteria. We therefore hypothesize that, by analogy with eukaryotes, an internal cytoskeleton maintains *Spiroplasma* membrane morphology.

**Figure 1 -.**
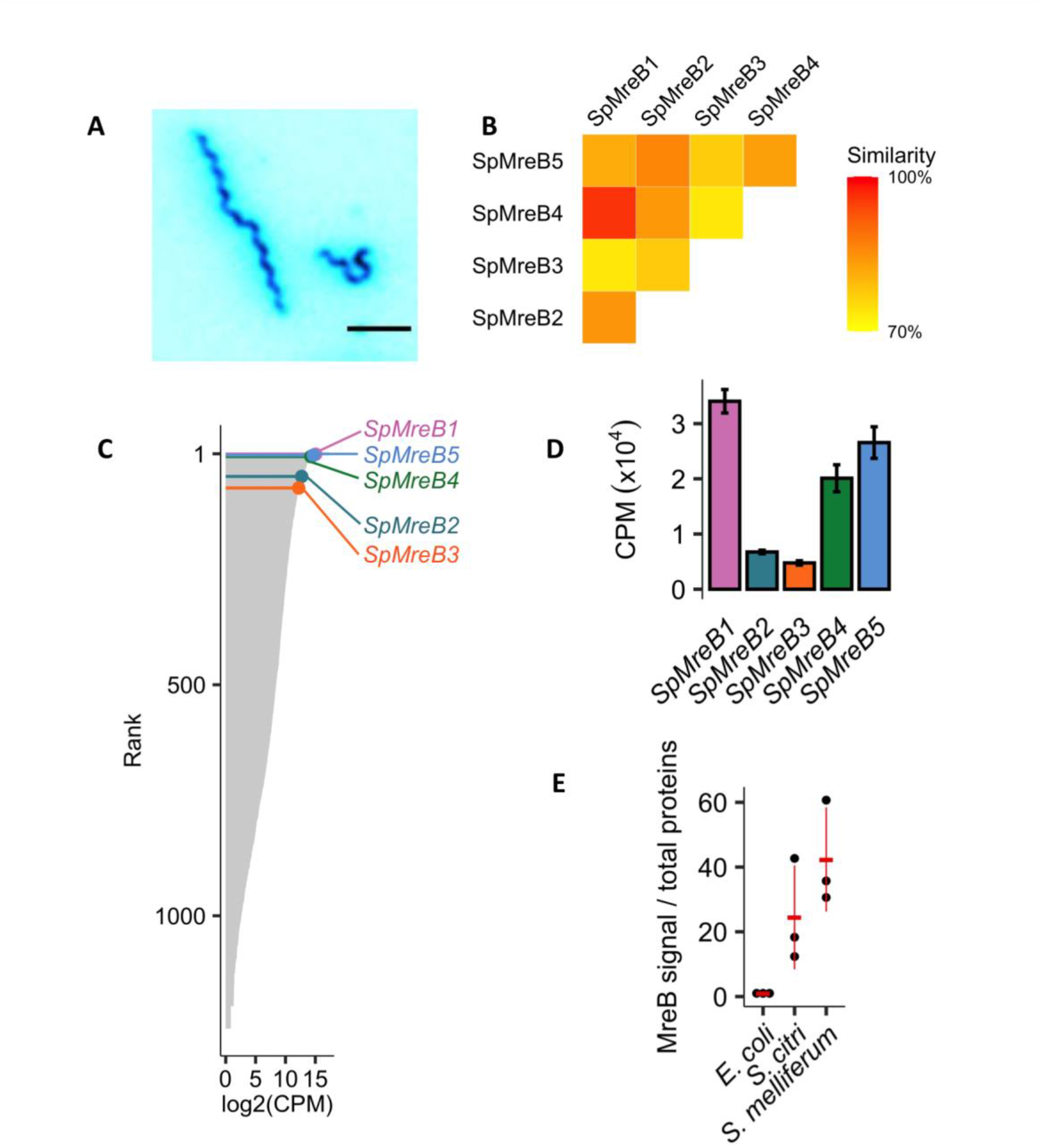
*Spiroplasma* MreBs are all strongly expressed *in vivo*. (A) Representative picture of *Spiroplasma poulsonii* grown *in vitro*. Scale bar = 5 μm. (B) Matrix of the percentage of similarity between SpMreBs protein sequences. Similarity was calculated using Geneious proprietary alignment algorithm and BLOSUM45 scoring matrix. (C) Relative expression of *SpMreB*s compared to *S. poulsonii* transcriptome. Grey bars indicate all 1491 transcripts ranked from the most to the least expressed. Lollipops indicate the position of each *SpMreBs*, in the top 5% most expressed genes. (D) Expression level of each isoform, in Counts Per Million (CPM) from (48). (E) Western blot quantification of MreB in *E. coli, S. citri* and *S. melliferum*. MreB signal was normalized to the total amount of proteins on each track as revealed by Coomassie blue staining. *Spiroplasma r*atios were normalized to 1 on the *E. coli* ratio. Each black dot represents the quantification on an independent biological replicate; the red bars and segments indicate the mean and standard deviation, respectively.

Early electron cryotomograms of *Spiroplasma* have revealed intracellular filament that may function as a membrane-associated cytoskeleton (14). One candidate protein called Fibril (Fib), whose polymerization forms an internal ribbon spanning the entire length of the bacterium, may provide structural support for the membrane (12, 14–18). Therefore, Fib was initially thought to be necessary and sufficient to maintain the helical shape of *Spiroplasma*. Yet, the serendipitous isolation of a non-helical *Spiroplasma* variant still harboring fibrils indicates that Fib is not sufficient to maintain helical shape (19). A proteomic analysis of purified ribbons confirmed the massive presence of Fib, but also revealed the presence of MreB, a bacterial homologue of the eukaryotic actin (17, 20, 21), indicating it could play a role in maintining *Spiroplasma’s* helical shape.

MreB plays an essential function in determining the shape of almost all non-spherical bacteria (4, 22). MreB polymerizes in an ATP-dependent fashion to form anti-parallel double filaments that interact with the cytoplasmic tail of the transmembrane protein RodZ (23–25). The periplasmic part of RodZ then positions the cell wall synthesis machinery, thus linking the cell wall expansion sites (hence cell elongation) to the MreB subcellular localization (26). Early studies indicated that MreB forms filaments spanning the cell length (27, 28), crystallizing a hypothesis wherein MreB is a cytoskeletal protein, providing mechanical support to the cell envelope. Despite mechanical characterization that was consistent with this hypothesis, subsequent experiments suspected these filaments were artifactual (23, 29, 30). Fluorescent protein fusions with improved functionality and limited artifacts have helped demonstrate that MreB forms short polymers in close interaction with the membrane. These filaments undergo a processive circumferential movement around the cell as they coordinate the synthesis of new peptidoglycan (31–35). In *E. coli*, MreB forms ~100 nm-long filaments, too short to mechanically support the cell envelope and constitute a cell wall. Helical filaments were however observed in other fusions in *B. subtilis* and *E. coli*, reviving the helix model (35, 36). These conflicting localization data could however be partially explained by variations between strains or observation protocols (35). In summary, MreB polymers maintain cell shape by patterning the cell wall, but not by directly bearing load as would be expected for a cytoskeleton.

Consistent with MreB’s function in cell wall patterning, *non-Spiroplasma* members of the wall-less Mollicutes family lost *mreB* (10). In striking contrast, *Spiroplasma* not only retained but multiplicated *mreB* to reach five to eight copies per genome depending on the species (37, 38). This suggests that the protein has a major alternative function in *Spiroplasma* physiology, evidently independent of cell wall synthesis (10, 38). A breakthrough in understanding *Spiroplasma* MreBs function has been achieved in the plant pathogenic species *Spiroplasma citri* (37). Sequencing of the naturally non-helical strain ASP-I isolated in 1980 revealed a truncation in the mreB5 gene, while *fib* and other *mreB* isoforms where intact (19). MreB5 forms short filaments *in vitro*, can interact both with the major cytoskeleton protein Fib and with liposomes. *mreB5* deletion mutants lose their helical shape and consequently their motility (37). These observations are consistent with a scenario where MreB5 participates in maintaining *Spiroplasma* shape. The mechanism by which MreB5 maintains shape and the functions for the other four *mreB* paralogues remain however still unknown.

Here we investigate *Spiroplasma* MreBs functions and interactions using *Spiroplasma poulsonii*, a close relative of *S. citri* that is a natural endosymbiont of *Drosophila melanogaster* (39). *S. poulsonii* lives in the fly hemolymph (a functional equivalent of mammal blood) and gets vertically transmitted over generations by infecting oocytes during oogenesis (40). *S. poulsonii* causes a male-killing phenotype whereby all male embryos are killed by the action of a secreted toxin, hence biasing the population towards females (41). *S. poulsonii* is an interesting model to investigate the selective effect of bacterial shape because of its rapid evolution and selection through vertical transmission, which points to strong selective pressure for helical shape *in vivo* (42). *Spiroplasma* in general are however poorly tractable bacteria *in vitro* (43, 44). Although some gene inactivation attempts were successful in *S. citri* (e.g. 5, 6), this taxon is characterized by an extremely inefficient recombination machinery due to the pseudogenization of *recA* (47–49), which is a major hurdle for genetic engineering. *S. poulsonii* especially has only been recently cultured *in vitro* (49) and transformed to express a fluorescent marker (50), but no genomic modification has been achieved so far in this species, thus preventing a systematic knock-out approach.

To explore the function of MreBs in *S. poulsonii*, we setup a heterologous expression of *Spiroplasma* MreB coding genes in *E. coli*. By individually expressing the three isoforms, we showed that each is able to polymerize in its own filamentous pattern. By systematically co-expressing isoforms in all possible combinations, we found that they together form a complex network of interactions regulating each other’s polymerization patterns, which potentially determines MreBs assembly in *Spiroplasma*. In the light of these results, we propose a model of MreBs interactions and discuss why five different isoforms would be necessary to maintain morphology and motility in *Spiroplasma*.

## Results

### *Spiroplasma* strongly expresses MreBs

*S. poulsonii* possesses five *mreB* paralogues distributed on three chromosomal loci (38, 49). *S. poulsonii* MreBs (SpMreBs) have a remarkably high level of sequence similarity one with another (*i.e*. an identical amino-acid or an amino-acid with similar chemical properties at a given position), ranging between 73% for the SpMreB1-SpMreB3 comparison and 96% for the SpMreB1-SpMreB4 comparison (Figure 1B). We inspected the transcriptome of S. poulosnii in axenic liquid cultures for the expression levels of MreB. In these conditions, *S. poulsonii* maintains its characteristic helical shape as the one it exhibits *in vivo*. Transcriptomics data indicate that all five isoforms are expressed at high level, ranking among the top 5% most expressed genes in *S. poulsonii* (Figure 1C) (49). Among them, *SpMreB1, 4* and *5* have a higher expression level than *SpMreB2* and *3* (Figure 1D). This suggests that SpMreB1, 4 and 5 are more abundant than 2 and 3. We thus explored MreB protein levels in Spiroplasmas. While we could not confirm this for *S. poulsonii* due to the lack of cross-reactivity of anti-MreB antibodies, Western blots on proteins from *S. citri* and *S. melliferum*, two other *Spiroplasma* species, show that the ratio of MreB-specific signal over total proteins was 20 to 40-fold higher than in *E. coli* (Figure 1E). This demonstrates that MreB represents a massive part of the *Spiroplasma* proteome compared to that of *E. coli*. Our transcriptome data along with the relative abundance of MreB in related *Spiroplasma* species altogether indicate that all SpMreB isoforms are strongly expressed in *S. poulsonii*.

### SpMreBs form long filaments

The high abundance of MreBs in *Spiroplasma* suggests they play an architectural function in cells. In addition, functional domains such as intra- and inter-protofilament binding domains and the ATP-catalytic pocket are conserved between SpMreBs and MreBs from rod-shape bacteria, which suggests that SpMreBs can form polymers (Figure S1) (38). We hypothesize that SpMreB concentration plays a role the formation of intracellular cytoskeletal structural that could participate in shaping *Spiroplasma* cells.

We therefore investigated the polymerization behavior of individual SpMreBs. As expected from their evolutionary history (37), SpMreB4 and SpMreB5 are highly similar to SpMreB1 and SpMreB2 (96% and 86.4%, respectively). We therefore focused on SpMreB1, −2 and −3. We circumvented the weak genetic tractability of *Spiroplasmas* by resorting to the heterologous expression of fluorescently tagged SpMreB in *Escherichia coli* cells. Heterologous expression has been instrumental in investigation of MreB structure and dynamics (51, 52). Here, we built a sandwich fusion where the coding sequence for a monomeric superfolder Green Fluorescent Protein (GFP) is inserted on a poorly conserved external loop, generating a functional MreB fusion with limited alterations of its native structure (25, 53–55). The sandwich fusion to the native *E. coli* MreB (EcMreB^GFP^) displayed the characteristic peripheral diffraction-limited puncta pattern along the membrane (53) (Figure 2A). In contrast, the three SpMreB^GFP^ isoforms formed distinctive filament-like structures when expressed at high levels as in *Spiroplasmas*. SpMreB1^GFP^ formed filaments a few micrometer-long extending along the cell. SpMreB2^GFP^ formed long and thick filaments along the cell, which connected large puncta. SpMreB3 mainly formed transverse filaments forming ring-like structures across the cell (Figure 2A).

**Figure 2 -.**
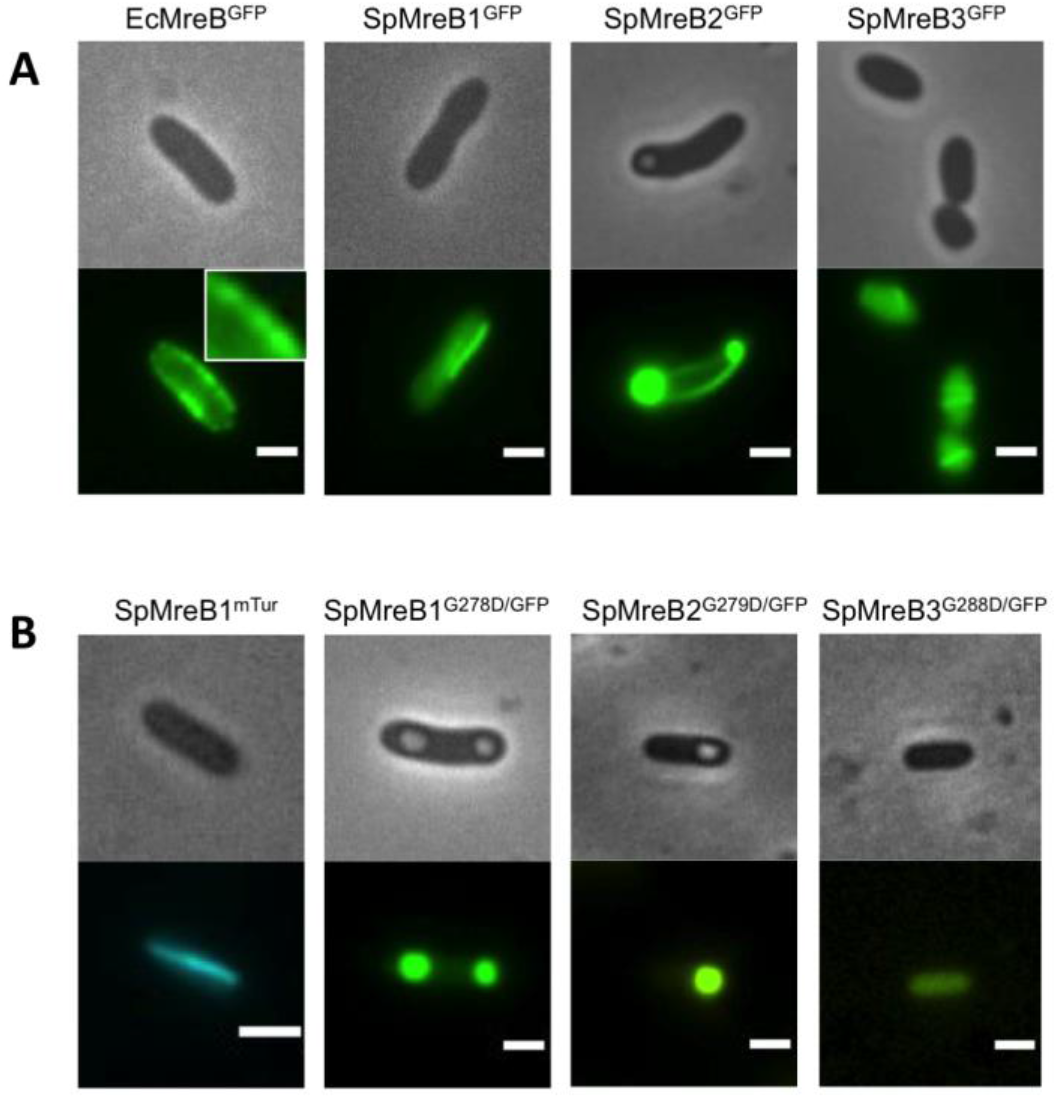
SpMreBs forms filaments when expressed in *E. coli* cells. (A) *E. coli* cell expressing MreB^GFP^ constructs. The typical peripheral puncta along the membrane are observed with EcMreB^GFP^ (insert is a 2X magnification), SpMreB1^GFP^ shows longitudinal filaments; SpMreB2^GFP^ shows pole puncta and longitudinal filament; SpMreB3^GFP^ shows transversal filaments. (B) *E. coli* cells expression SpMreB1^mTur2^ showing similar longitudinal filaments to that formed by SpMreB1^GFP^; or expressing the point-mutated constructs SpMreB1^G278D/GFP^, SpMreB2^G279D/GFP^ and SpMreB3^G288D/GFP^ (mutation on the intra-filament interaction region). Scale bar = 2 μm.

MreB fusions are prone to artifactual polymerization, in particular at high expression levels (53). However, compounds such as A22 (56) and MP265 (57) that inhibit MreB polymerization in other species causing loss of native morphology were ineffective on SpMreBs, preventing their use in validating SpMreBs polymer formation. To control for potential artifacts, we instead generated SpMreBs point-mutants in the intra-filament interaction region and visualize polymerization phenotypes (23, 38) (Figure S1). None of the SpMreB1^G278D/GFP^, SpMreB2^G279D/GFP^ and SpMreB3^G288D/GFP^ mutants formed filaments. Instead, these mutants showed either accumulated signal in puncta or displayed no signal (Figure 2B), indicating that wild-type constructs formed polymers that are unlikely to be artifactual. Consistent with this, we could observe filaments similar to SpMreB1^GFP^ when swapping GFP to the fluorescent protein mTurquoise2 (Figure 2B), further confirming that the tag does not affect the filament formation. Finally, we visualized native EcMreB in SpMreB-expressing *E. coli* by immunofluorescence. We could not observe any misshaping or change in localization pattern of EcMreB, which indicates that there are no confounding interactions between SpMreBs and the native EcMreB (Supplementary Information Text and Figure S2A-F).

We then looked into the mechanism of filament formation. We hypothesized that protein-protein interactions between SpMreBs drive the formation of polymers. To test this hypothesis, we measured the interaction of each isoform with itself using a two-hybrid system based on the reconstitution of the adenylate cyclase of *E. coli* upon protein-protein interaction (58) (Figure S2G). We observed a strong BACTH signal for all isoforms indicating that each SpMreB interact with themselves, which likely leads to polymerization and filament formation observed in Figure 2A.

### *Spiroplasma* MreB filaments are polymorphic and dynamic

Remarkably, while fluorescent EcMreB gave a consistent pattern from one cell to another, SpMreB^GFP^ expressing cells had an important phenotypic heterogeneity. We thus carefully categorized the morphology of the filaments formed by single-isoform SpMreB^GFP^ in the heterologous expression system. Most cells formed filaments (Figure 3A). Quantitative analysis revealed that the filament patterns shown in Figure 2A are isoform-specific: longitudinal filaments are proper to SpMreB1^GFP^, puncta associated with filaments proper to SpMreB2^GFP^ and transversal filaments proper to SpMreB3^GFP^ (Figure 3A). We found that for all isoforms, only a small proportion of cells had a homogenously diffuse cytoplasmic signal. Also, only a small proportion of cells displayed puncta without filaments in SpMreB2 and 3.

**Figure 3 -.**
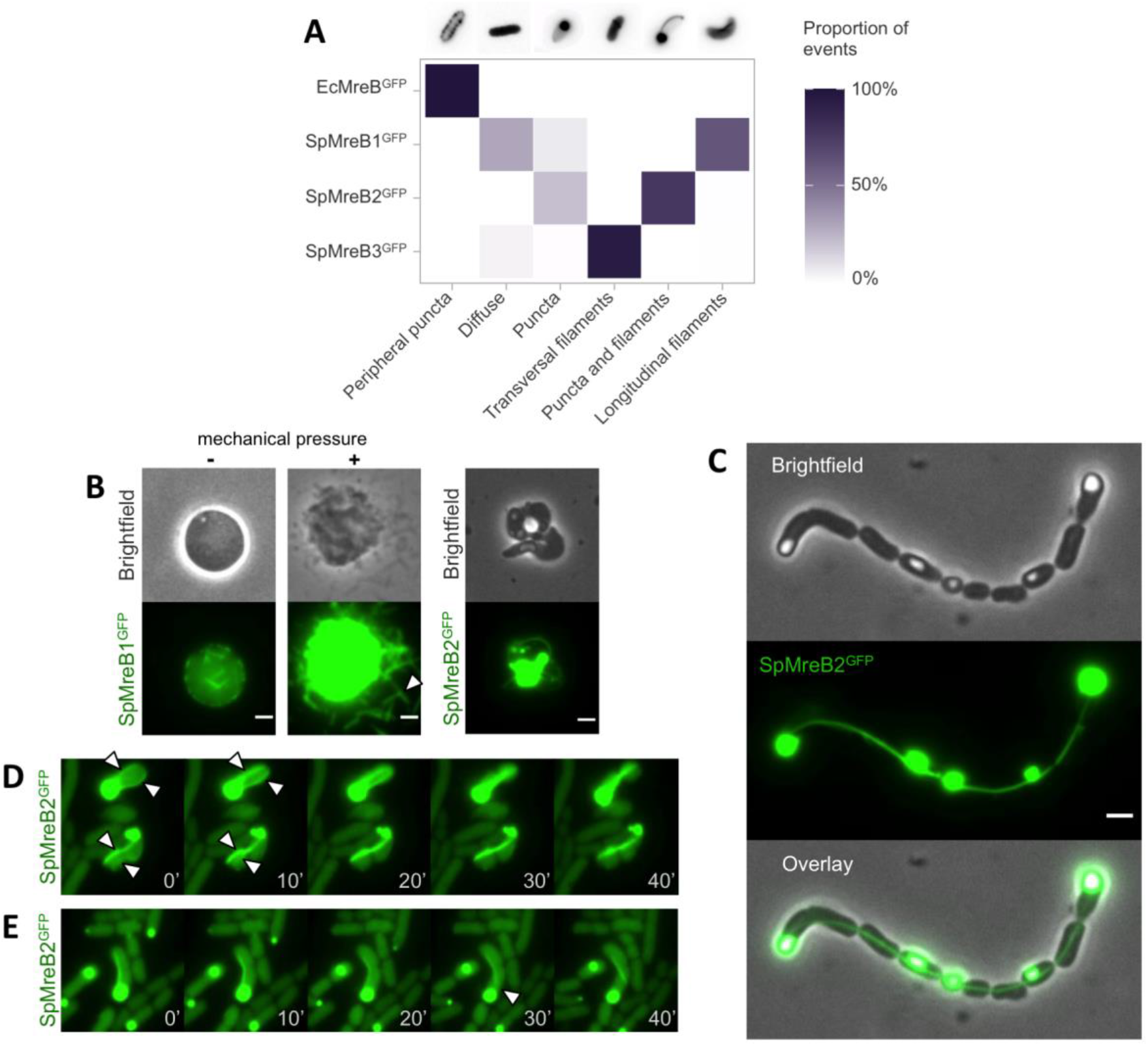
SpMreBs have distinct filament pattern and dynamics. (A) Identification of the main filament patterns and quantification of their observed proportion upon expression of single MreB^GFP^ construct. Color density indicates a percentage of observations for each condition, from n > 250 observations from three independent replicates. (B) SpMreB^GFP^ upon strong induction (1 mM IPTG) showing cell rounding for SpMreB1^GFP^ and release of the filaments in the close environment upon mechanical cell lysis; and cell misshaping for SpMreB2^GFP^. (C) Overexpression of SpMreB2^GFP^ resulting in a string of cells connected by a thick filament. (D) Timelapse of SpMreB2^GFP^ proto-filament fusion. White arrows show the proto-filaments before their merging. (E) Timelapse of a SpMreB2^GFP^ filament detaching from a punctum. The white arrow indicates the moment and location when the filament detaches. Times from the first pictures are indicated on each sub-panel in minutes. Scale bars = 2 μm.

Increasing the induction level of the SpMreB1^GFP^ construct to 1 mM IPTG (a concentration commonly used for recombinant protein production) lead to swollen, round cells. These easily rupture upon mechanical pressure and release filaments in the vicinity of the dead cell (Figure 3B), suggesting that they are not tightly attached to the cell envelope. Overexpressing SpMreB3 did not yield any interpretable observations as most of the surviving population seemed to systematically lose fluorescence. Strongly inducing SpMreB2 with 1 mM IPTG resulted in cells with consistently aberrant shapes, accompanied with puncta accumulation and swelling (Figure 3B). We occasionally observed strings of normally-shaped cells connected by a continuous SpMreB2 filament (Figure 3C). This could be the result of the inability of *E. coli* to cut the filament transversally during division; hence the structure is extending from between two daughters, blocking their separation.

EcMreb filament processivity drives a net movement of filaments along the cell. We therefore wondered whether SpMreB were also mobile. We thus performed dynamics visualization of if the SpMreB1, −2 and −3 filaments by timelapse microscopy in our heterologous system. Visualizations of SpMreB1 and SpMreB3 filaments over two hours did resolve any mobility during or after polymerization. In contrast, SpMreB2 filaments were dynamic after completion (Figure 3D and 3E and Supplementary Movies 2 and 3). Specifically, we observed multiple filaments undergoing lateral displacements. When cells had several filaments, they merged into thicker bundles, generally associated along the cell envelope (Figure 3D and Supplementary Movie 2). These data allowed us to estimate that SpMreB2 protofilaments move at a rather slow speed ranging from 30 to 60 nm/min. For comparison, EcMreB filaments move at 1 to 5 μm/min (36).

### SpMreBs form an interaction network *in vivo*

Cryo-electron tomography performed on *Spiroplasma melliferum* cells have provided an empirical basis for the ultrastructure of the *Spiroplasma* cytoskeleton. Two models were inferred from these tomograms. The first model (three-ribbons model) proposes that the cytoskeleton is composed of a ribbon of thin filaments (presumably formed by MreBs) sandwiched between two ribbons of thicker filaments (presumably formed by Fibril proteins) (14). Alternatively, the second model (one-ribbon model) proposes that a single ribbon composed of mingled Fib and MreBs forms the cytoskeleton (17). In both models however, the cytoskeleton is only composed of one or two filament types. Therefore, the five polymer-forming MreBs and Fib that compose the filaments are very likely to interact with one another.

To identify the potential interactions between SpMreB isoforms, we first employed a high throughput approach and performed co-immunoprecipitation (co-IP). We overexpressed and purified single SpMreBs^GFP^ and incubated them with a total protein extract of wild-type *S. poulsonii*. Complexes formed between the SpMreBs^GFP^ fusion proteins and native *S. poulsonii* were then purified, broken down and their components were analyzed by mass spectrometry (LC-MS/MS). This approach identified 231 *S. poulsonii* proteins interacting with at least one SpMreB isoform. Remarkably, each isoform had at least one other isoform amongst its interactors, indicating that they most likely all act in combination i*n vivo* to build the *Spiroplasma* cytoskeleton (Figure S3). We thus identified the direct interaction network between isoforms (Dataset S1 and Figure S3). Only SpMreB1 interacts with Fib, suggesting that it could serve in anchoring the SpMreBs structure to the Fib ribbon. As this co-IP analysis suggests *in vivo* interactions between isoforms, we went on to test specific interactions between each isoform and their effect on the regulation of polymerization.

### The SpMreBs interaction network orchestrates polymerization

We then investigated whether the inter-SpMreBs interactions identified by co-IP could regulate their polymerization and filament formation. We thus simultaneously expressed several isoforms in *E. coli* and visualized the resulting polymerization patterns, which we could then compare to single isoform expression (Figure 2). We first co-expressed SpMreB1, −2 and −3 each tagged with a distinct fluorescent protein (SpMreB1^GFP^, SpMreB2^mTurquoise2^ and SpMreB3^mScarlet^) under the control of a single inducible promoter. All tags could be detected in the cell population. Single filaments in most cells were composed of the three isoforms, suggesting that they are able to form either mixed polymer or assembled homopolymers (Figure 4A). Protein expression and filament morphology were however more heterogeneous between cells, compared to individual expressions.

**Figure 4 -.**
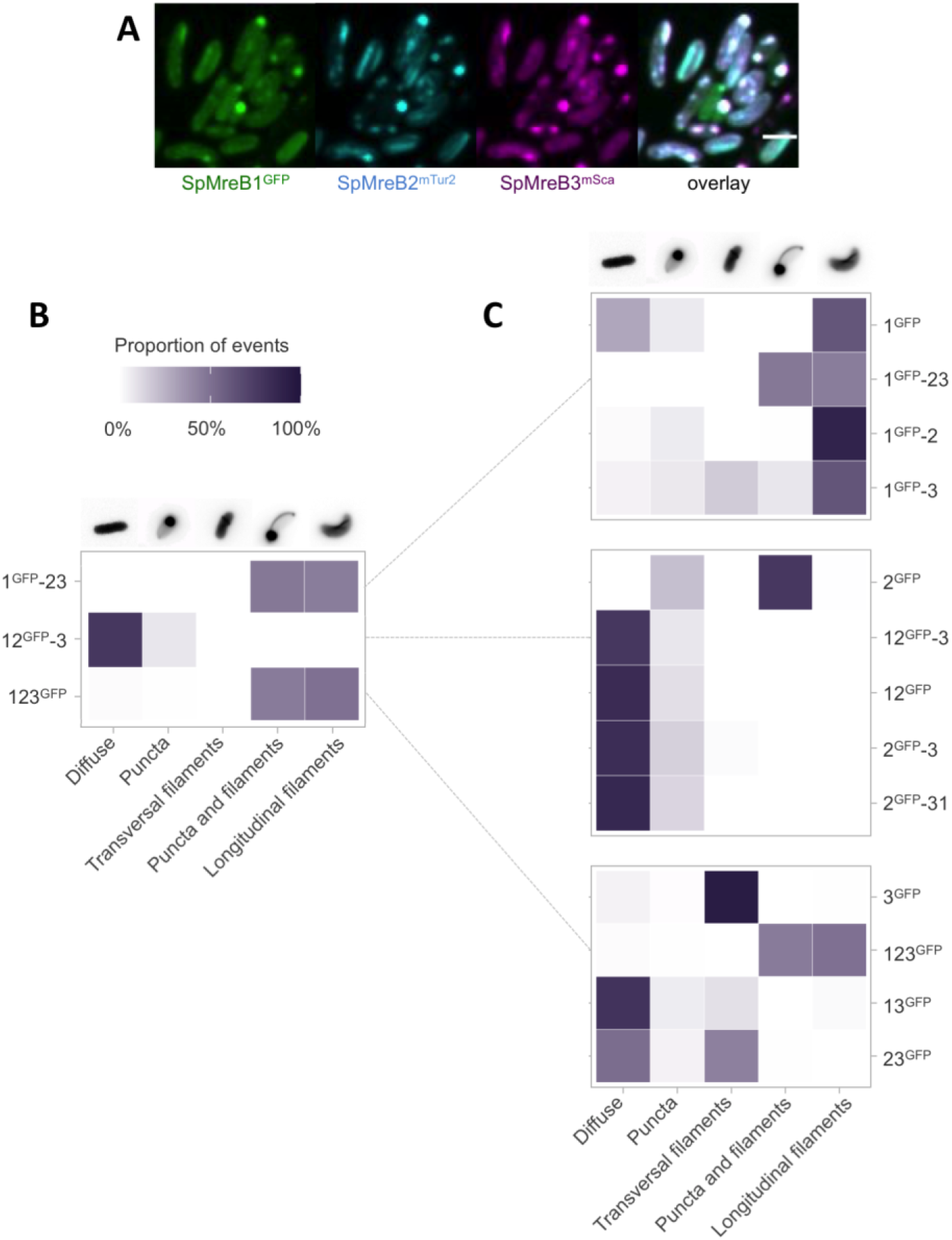
SpMreB interactions regulate polymerization. (A) Overlay of *E. coli* cells expressing pET28-SpMreB1^GFP^-SpMreB2^mTurq2^-SpMreB3^mScarlet^. Scale bar = 2 μm. (B and C) Identification of the main patterns and quantification of their proportion upon expression of transcriptional fusion construct with the GFP tag on SpMreB1, SpMreB2 or SpMreB3. Color density indicates a percentage of observations for each condition, from n > 250 observations from three independent replicates.

To rigorously delineate the function for the SpMreB interaction network, we therefore used a combinatorial approach based on the observation of a single-tagged isoform co-expressed with untagged isoforms. For simplicity, we named these combinations using only their isoform numbers with a superscript “GFP” directly following the number of the tagged isoform (*e.g*. 1^GFP^-23 refers to the construct SpMreB1^GFP^-SpMreB2-SpMreB3; see Figure S4). If all isoforms are part of a single structure, we should observe similar fluorescence patterns independently of the GFP tag position. This was the case for SpMreB1 and SpMreB3 as indicated by the identical pattern of 1^GFP^-23 and 123^GFP,^ for which we both observed cells with longitudinal filaments and cells with puncta and longitudinal filaments in similar proportions (Figure 4B). SpMreB2 in 12^GFP^-3 however displayed a diffuse phenotype, distinct from the other two combinations, and also different from its single expression. This confirms that SpMreB filaments do not simply form a single structure but rather act through an elaborate interaction network regulating filament formation.

To further characterize how interactions between isoforms regulate filament formation, we turned to co-expression of SpMreB pairs. The proportion of longitudinal filaments observed in the single 1^GFP^ expression was lower in 1^GFP-^3 and 1^GFP-^23, but remained at single expression levels in in 1^GFP-^2. This indicates that SpMreB3 modulates SpMreB1 polymerization (Figure 4C) while SpMreB2 has little to no effect on SpMreB1. Coexpression where the GFP tag is on SpMreB3 gave a more complex phenotype. Transversal filaments were observed in almost all cells upon SpMreB3^GFP^ single expression (Figure 2A and 3A), while none of these filaments were detected with the 123^GFP^ construct (Figure 4B). This indicates that the presence of both SpMreB1 and SpMreB2 inhibits SpMreB3 transversal filament formation. However, such transversal filaments were observed with both the 13^GFP^ and 23^GFP^ constructs (Figure 4C), indicating that SpMreB1 and SpMreB2 must be both present to completely inhibit their formation. The proportion of cells harboring transversal filaments was higher upon 23^GFP^ expression (44% on average) than 13^GFP^ expression (10% on average), indicating a stronger inhibition from SpMreB1 than from SpMreB2. Collectively, this indicates that SpMreB1 and SpMreB3 form filaments together while having a limiting effect on each other’s polymerization.

12^GFP-^3 on the other hand had its proper pattern with a majority of cells displaying a diffuse cytoplasmic signal and a few harboring puncta (Figure 4B). This phenotype is independent of the order of transcription of the isoforms which may affect stoichiometry, as it is identical to that observed with a 2^GFP^-31 construct (Figure 4C) (59). This suggests that SpMreB1 and/or SpMreB3 inhibit SpMreB2 filament formation. Both 2^GFP-^3 and 12^GFP^ displayed similar results to 12^GFP-^3, indicating that the presence of any of the other isoforms can inhibit SpMreB2 filament structuring (Figure 4C).

## Discussion

The conservation and duplications of MreB coding genes in *Spiroplasma* raises major questions regarding their function in wall less bacteria (10, 38). Here we report evidence supporting that all SpMreBs isoforms are able to form filaments *in vivo* and that each isoform can affect the polymerization pattern of others through a complex network of interactions. As a consequence, this suggests that [SpMreB1-SpMreB4], [SpMreB2-SpMreB5] and [SpMreB3] clusters have distinct functions and are not necessarily redundant (Figure S5). From our observations, we can infer the existence of at least two separate structures *in vivo*, one involving SpMreB1 (and possibly the closest homologue, SpMreB4) with SpMreB3 and one involving SpMreB2 (and possibly the closest homologue, SpMreB5). Yet we uncovered interaction between almost all isoforms in the co-immunoprecipitation experiment, suggesting that although polymeric structures are distinct, the monomers themselves are probably able to interact one with another.

SpMreB1 formed static filament structures that do not attach to the cell envelope (Figure 3D). We thus hypothesize that it could form a backbone structure on which other isoforms would associate. Its interaction with Fib suggests a potential association with the Fib ribbon, hence coordinating Fib and MreB functions. SpMreB1 function is also interesting in the perspective of its interaction with SpMreB3, which likely integrates in a common structure (Figure 4B). Interestingly, SpMreB3 was the only isoform producing transversal filaments, similar to the orientation of EcMreB polymers. Furthermore, an amphipathic helix has been predicted on the N-terminus region of *Spiroplasma* MreB3s (38), suggesting an ability to attach to membranes. The *S. citri* ASP-I mutant that has intact MreB1 and MreB3 isoforms is not helical, which indicates that SpMreB3 is not sufficient to form helical cells (37). SpMreB3 could instead serve as a set attachment point between SpMreB1 filaments and the membrane and twist them to follow the cell body helicity maintained by SpMreB5 (and possibly SpMreB2).

*Spiroplasma* swims by changing shape. Cells deform their membrane by producing a kink in their helix. The propagation of this kink along the cell body propels the cell forward. Kinks are produced by a local and processive change in the cell body helicity (13), but the exact mechanism regulating helicity and handedness is still elusive. We suspect that active motion of MreB and Fib filaments relative to each other can produce these deformations. We found that SpMreB2, the closest homologue *S. citri* MreB5 (37), was the only isoform producing mobile filaments. A proposed model based on a single-ribbon cytoskeleton involves a coordinated length change of the filaments forming the internal ribbon, with unknown proteins anchoring the moving fibrils to the membrane to propagate movement to the cell (12, 60). In the three-ribbons cytoskeletal model, the shortening of one ribbon would be sufficient to cause a change in the helix handedness, resulting in the formation of the kink (14). In this case, only one ribbon need to be mobile while others could stay fixed and serve as an anchoring point. SpMreB2 polymerization pattern involved large puncta and filaments that are connected to the puncta but are also attached to the cell body on their own (see Figure 3A and Figure 3C). The puncta themselves could serve as attachment structures as their localization seems to be defined at the cell pole or more rarely at the cell center. SpMreB2 filaments could directly attach to the membrane since the homologous MreB5 can interact with liposomes (37). Taken together, the mobility and attachment features of SpMreB2 make it a promising candidate as a regulator of *Spiroplasma* motility. An attractive hypothesis would be that [SpMreB1-SpMreB3] (and possibly SpMreB4) form fixed structures anchored to the membrane (by SpMreB3 and the Fib ribbon (by SpMreB1), against which the [SpMreB2-SpMreB5] structure slides to produce and propagate the kinks.

Last, the SpMreB2 filaments grew into long and mechanically robust filaments upon over-expression of the single construct that are deleterious to *E. coli* growth. We hypothesize that a *Spiroplasma*-specific mechanism regulates SpMreB2 filament length in native conditions. This mechanism could involve a yet uncharacterized protein capable of cutting the filament or limiting its growth (possibly another SpMreB isoform). Alternatively, the filament could be split longitudinally during cell division (yielding back the thinner protofilaments observed before merging), which would be in accordance with the remarkable division process of the bacterium whereby cells split longitudinally (61).

Collectively, our results indicate that the sub-functionalization of each isoform and their interactions one with another allows building a complex polymeric inner structure. Despite rigorous mechanical characterization experiments that highlight a load-bearing function of MreB in *E. coli* (62), the hypothesis where MreB filaments provide mechanical stability has been largely abandoned, favoring a model where MreB almost exclusively functions by patterning the cell wall. The high abundance of SpMreB isoforms *in vivo*, as well as the high induction level required to observe filaments in our heterologous system, is rather reminiscent of eukaryotic actin. To form a robust cytoskeleton, mammalian cells must maintain a high concentration of intracellular actin monomer to maintain polymerization, making actin one of the most highly expressed protein in a cell. In the light of our data and of *Spiroplasma* biology, we propose that the MreB polymeric structure coordinates the Fib cytoskeleton and the membrane to maintain *Spiroplasma* helicity and generate kink-movement. The example of *Spiroplasma* MreBs highlights the divergent evolution of these proteins in wall-less bacteria, where they control bacterial shape through mechanisms that are independent of peptidoglycan synthesis.

## Material and Methods

### Sequence analysis and alignment

MreB coding sequences were retrieved from the *S. poulsonii* reference genome (Accession number JTLV02000000) and aligned using Geneious 11.0.5 (https://www.geneious.com). Similarity was calculated on the translated nucleotide sequences based on a BLOSUM45 scoring matrix. Functional sites were predicted based on previously published sequence analyses on MreBs from other *Spiroplasma* species (38).

### Bacterial strains and plasmids

*E. coli* and *S. poulsonii* strains used in this work are listed in Supplemental Table 1. As *Spiroplasma* have an alternative genetic code, coding sequences for MreB isoforms were codon-optimized and fully synthesized by Genewiz (South Plainfield, NJ, USA). Two constructs for each isoform, each with a different codon optimization, were ordered and indifferently used in constructs after controlling that codon optimization was not affecting the polymerization patterns. Plasmid construction was made by or by restriction/ligation or by Gibson Assembly, using primers listed in Supplemental Table 2. Constructs were built in competent XL10-Gold cells (Agilent) and subsequently transformed in the appropriate strain for experiments.

### Live fluorescence microcopy

*E. coli* was grown in LB with 50 μg/mL kanamycin for 16 to 18 hours at room temperature (22-23°C) under 300 rpm shaking. Induction was made using 100 μM IPTG for SpMreB constructs and 50 μM IPTG for EcMreB constructs, unless otherwise stated. *S. poulsonii* was extracted from *Drosophila* flies and grown in BSK-H-spiro medium at 25°C with 5%CO2 and 10% O2, as previously described (49). Bacteria were observed on glass-bottom dished with a thin agarose pad on top. Images were acquired on a Nikon Eclipse Ti-E microscope. Constructs with three different fluorescent tags were observed on a Nikon Eclipse Ti2 microscope coupled with a Yokogawa CSU W2 confocal spinning disk unit. Mechanical pressure to break cells consisted in pressing the agarose pad with the finger.

### Filament quantitative analysis

The extremely high diversity of patterns that we uncovered with tagged SpMreB made already published segmentation methods inefficient, as well as machine-learning based image analysis. Images were thus manually screened and bacteria were classified according to the major pattern observed. Bacteria with no clear pattern were classified as “others” and not accounted for in the figures (<1% of observations). Bacteria were randomly picked from at least two fields of view per replicate, and three independent replicates per construct, until reaching a minimum of 300 observations per construct.

### Immunofluorescence microcopy

Bacteria were grown in LB with 50 μg/mL kanamycin and 100 μM IPTG for 16 to 18 hours at room temperature (22-23°C) under 300 rpm shaking. Cells were fixed in growth medium with 1.6% formaldehyde and 0.01% glutaraldehyde for 1 h at room temperature, and immunostained as previously described (63). EcMreB was detected using a polyclonal rat anti-*Bacillus subtilis* MreB (1:300) (27) that cross-reacts with that of *E. coli* but not *S. poulsonii*, and a secondary anti-rat antibody coupled with Alexa Fluor 488 (1:1000). Cells were observed on a Nikon Eclipse Ti-E microscope using 480 nm excitation and 535 nm emission filter sets.

### Western Blot

Wild-type *E. coli* MG1655 were grown in LB for 16 to 18 hours at room temperature (22-23°C) under 300 rpm shaking. *Spiroplasma citri* and *Spiroplasma melliferum* were grown for 24 hours at 29°C without shaking in SP4 medium. Cells were washed three times in PBS before being resuspended in SDS-Tris-Glycine buffer and boiled at 95°C for 15 minutes. Total proteins were then separated by SDS-PAGE, transferred to a nitrocellulose membrane and blocked for 30 minutes with 2% BSA in PBS-Tween 0.1%. MreB was detected using a polyclonal rat anti-*Bacillus subtilis* MreB (1:3000) incubated overnight at 4°C and a secondary anti-rat antibody coupled with horseradish peroxydase (1:10000). Detection was performed with an ECL kit (Amersham). For normalization, an identical amount of proteins has been deposited on another SDS-PAGE run in parallel and stained with the Coomassie blue InstantBlue protein staining (Expedeon). The bioluminescence signal (MreB) was quantified using ImageJ built-in functions and normalized to the total protein amount approximated by the total InstantBlue signal.

## Supporting information

Supplementary Information Tex

## Acknowledgments

We are grateful to Rut Carballido-López, Yuko Inclan, Nikolay Ouzounov, Alice Cont, Handuo Shi and Kerwyn Casey Huang for providing critical reagents and plasmids. This work was supported in part using the resources and services of the Proteomics Research Core Facility at the School of Life Sciences of EPFL and we are especially grateful to Romain Hamelin and Florence Armand for their input on the co-immunoprecipitation experiment.

## Author contributions

FM and AP designed research; FM and XP performed research; FM analyzed data; AP and BL supervised the project; FM and AP wrote the paper. All authors contributed to the manuscript editing and validated the final version.

