## Supplementary Information Tex for "The wall-less bacterium *Spiroplasma poulsonii* builds a polymeric cytoskeleton composed of interacting MreB isoforms"

**This file includes:**

Supplementary information text

Supplementary methods

Figures S1 to S5

Tables S1 to S3

Legend for Dataset S1

Legend for Movie S1 to S3

SI References

**Other supplementary materials for this manuscript include the following:**

Dataset S1

Movie S1 to S3

Supplementary Information Text

**Interactions between EcMreB and SpMreBs**

We assessed the robustness of our heterologous expression approach by investigating the interaction of our constructs with the native MreB of *E. coli.* We showed that a ∆mreB mutant strain (1) cannot be complemented with an untagged version of SpMreB1, as expected (Fig. S2A). Previous studies showed that altering EcMreB polymerisation causes shape defects (2), we therefore co-expressed EcMreB and SpMreB1 in the ∆mreB strain. We did not observe any growth or shape defect upon co-expression with increasing levels of SpMreB1 (Fig. S2B-C), which suggests that it does not interfere with EcMreB. SpMreBs also had little effect on the cells morphology, although they appeared slightly elongated upon SpMreB1 and SpMreB2 expression and slightly thinner or wider upon expression of SpMreB1 or SpMreB2 and 3, respectively (Fig. S2D-E). Further, immunofluorescence on native EcMreB indicated that the peripheral puncta pattern was not disrupted upon expression of SpMreB1, -2 or -3 (Fig. S2F). Last, we tested if SpMreB isoforms can interact with EcMreB using the two-hybrid approach used to proof isoforms self-interactions. Here again, pairwise comparison of EcMreB with SpMreB1, 2 or 3 revealed no signal, further indicating that SpMreBs do not interact with EcMreB (Fig. S2G).

**Supplementary methods**

**Growth curves**

A Δ*mreB* strain with pRMmreBind-2 (1), a plasmid carrying a native copy of *mreB* under control of a pLac promoter, was transformed with a pTet-SpMreB1 carrying *SpMreB1* untagged under control of a pTet promoter. pRMmreBind-2 is induced by IPTG and repressed by glucose, while pTet-SpMreB1 is induced by anhydrotetracycline (ATc). Precultures from fresh colonies were grown LB with 100 µg/mL ampicillin and 50 µg/mL kanamycin for 8 hours at room temperature (22-23°C) under 300 rpm shaking, then diluted 1:10000 in growth medium with appropriate antibiotics and induced in 96 well-plates in triplicates. Plates were incubated under shaking at 25°C in a Tecan Infinite Pro 200 plate reader. OD_600_ was measured every 10 minutes for 20 hours.

**Morphology measurements**

Cell length and width were obtained from the brightfield channel of pictures acquired for filament quantification. Measurements were made using BacStalk software (3) with a segmentation cell size of 15 pixels, a minimum threshold of 7 pixels, and other parameters on their default value. Segmented images were manually examined to eliminate the segmentation of non-cell particles and bacteria that inactivated the transcript (no fluorescence of the GFP channel).

**Bacterial two-hybrid assay**

Two-hybrid assay was performed using the Bacterial Adenylate Cyclase-Based Two-Hybrid (BACTH) system (Euromedex) following the manufacturer’s instructions. Each *mreB* isoform was cloned into each BACTH plasmids using standard molecular biology methods. The constructs were cotransformed two by two in chemically competent *E. coli* BTH101. 5 μL of each transformation reaction was spotted on LB plates with 50 μg/mL kanamycin, 100 μg/mL ampicillin, 500 µM IPTG and 40 μg/mL X-gal. Plates were incubated at 30˚C for 48h prior to photography on a lightbox.

**Sample preparation for co-immunoprecipitation**

Bacterial cultures were performed as for microscopy experiments. 2 mL of overnight *E. coli* culture expressing single isoform with a GFP tag (or GFP alone as a negative control) were harvested, resuspended in 500 µL of Glucose-Tris-EDTA buffer (50mM glucose, 10 mM EDTA, 25 mM Tris-HCl pH 8) with 50 µg/mL of lysozyme and incubated 20 minutes at room temperature. DNAse I (1 µg/mL) and NP40 0.05% were then added to tubes, and samples were homogenized with 100 µm diameter glass beads on a Precellys Evolution (Bertin Technologies). Debris were pelleted for 1 minute at 10 000 g and the supernatant was used for the first incubation.

*S. poulsonii* was cultured as previously described (4)*.* 80 mL of 1 week-old culture was pelleted by 20 minutes centrifugation at 10 000 g at 18°C, washed twice with PBS 1.5X, and resuspended in 3 mL of lysis buffer (Tris 50 mM pH7.2, NaCL 300 mM, PMSF 1 mM, DNAse I 1 µg/mL, NP40 0,05%). Cells were homogenized with 100 µm diameter glass beads on a Precellys Evolution (Bertin Technologies) and incubated 20 minutes at room temperature, then for 2h on ice. Debris were pelleted for 1 minute at 10 000 g and the supernatant was used for the second incubation.

**Co-immunoprecipitation**

20 µL of GFP-Trap Magnetic Beads (Chromotek) were used for each sample, following the manufacturer’s instruction for preparation and magnet-washes. Beads were washed with 180 µL of PBS-Tween 20 (PBS-T) 0.1%, then saturated for 20 minutes with 2% BSA in PBS-T. 300 µL *E. coli* samples were then incubated for 1h at room temperature on a spinning wheel, and washed three times with PBS-T. 300 µL of *S. poulsonii* samples were then incubated on the beads for 1h30 at room temperature on a spinning wheel, and washed twice with PBS-T and once with mQ water. Elution was made by boiling the beads in 50 µL of Tris-Glycine SDS Running Buffer (Novex) for 5 min at 95°C. Each sample was made in triplicate. The protein concentration was assessed using the Pierce BCA Protein Assay Kit (Thermo Fisher Scientific) and 20 µg of sample was mixed with NuPage LDS buffer 1X final (Invitrogen) and 20 µM DTT.

**LC-MS/MS and data analysis**

LC-MS/MS and data analysis was performed at the Proteomics Core Facility of EPFL. Samples were separated by SDS-PAGE on a 10% polyacrylamide gel and stained with Coomassie blue. Each gel lane was entirely sliced and proteins were In-gel digested as previously described (5). Resulting peptides were desalted on StageTips (6) and dried under a vacuum concentrator. For LC-MS/MS analysis, resuspended peptides were separated by reversed phase chromatography on a Dionex Ultimate 3000 RSLC nano-UPLC system in-line connected to an Orbitrap Lumos Tribrid mass spectrometer (Thermo Fisher Scientific, Waltham, USA). Raw data were processed using MaxQuant 1.6.10.43 (7) against a concatenated database consisting of the Uniprot *Spiroplasma poulsonii* protein database (2003 entries LM201017), the Uniprot *Escherichia coli* protein database (4391 entries LM201019) and a list of MreB^GFP^ sequences used as baits. In order to reduce observed protein grouping artefacts resulting from the insertion of an internal GFP stretch into bait sequence constructs, the GFP stretch was manually removed. Carbamidomethylation was set as fixed modification, whereas oxidation (M), phosphorylation (S, T, Y), acetylation (Protein N-term) and glutamine to pyroglutamate were considered as variable modifications. A maximum of two missed cleavages were allowed and “Match between runs” option was enabled. A minimum of 2 peptides was required for protein identification and the false discovery rate (FDR) cutoff was set to 0.01 for both peptides and proteins. Label-free quantification and normalisation was performed by Maxquant using the MaxLFQ algorithm, with the standard settings (8).

The resulting data were processed using Perseus version 1.6.12.0 (9) from the MaxQuant tool suite. Reverse proteins, potential contaminants and proteins only identified by sites were filtered out as well as the protein groups from *E. coli* proteome. Protein groups containing at least two valid values in at least one group were conserved for the following analysis. Empty values were imputed with random numbers from a normal distribution (Width: 0.5 and down shift: 1.9 sd). A two-sample t-test with permutation-based FDR statistics (250 permutations, FDR = 0.05 or 0.01, S0 = 0.5) allowed determining significant candidates.

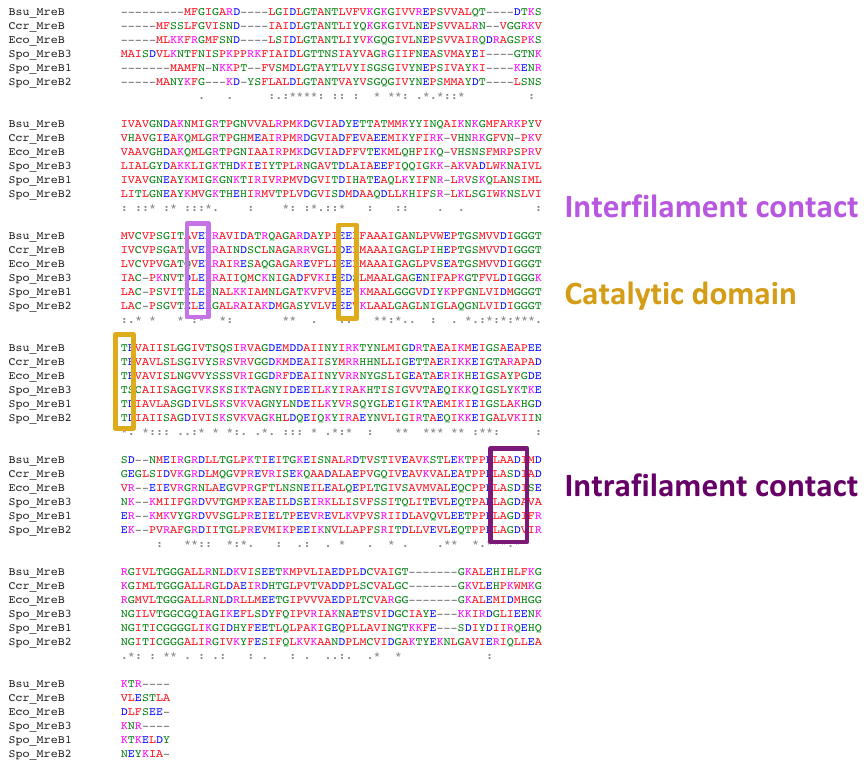

Fig. S1. Alignment of SpMreBs protein sequence with that of *E. coli* (Eco), *Bacillus subtilis* (Bsu) and *Caulobacter crescentus* (Ccr) showing the conservation of functional regions for inter- and intrafilament contact and ATP hydrolysis.

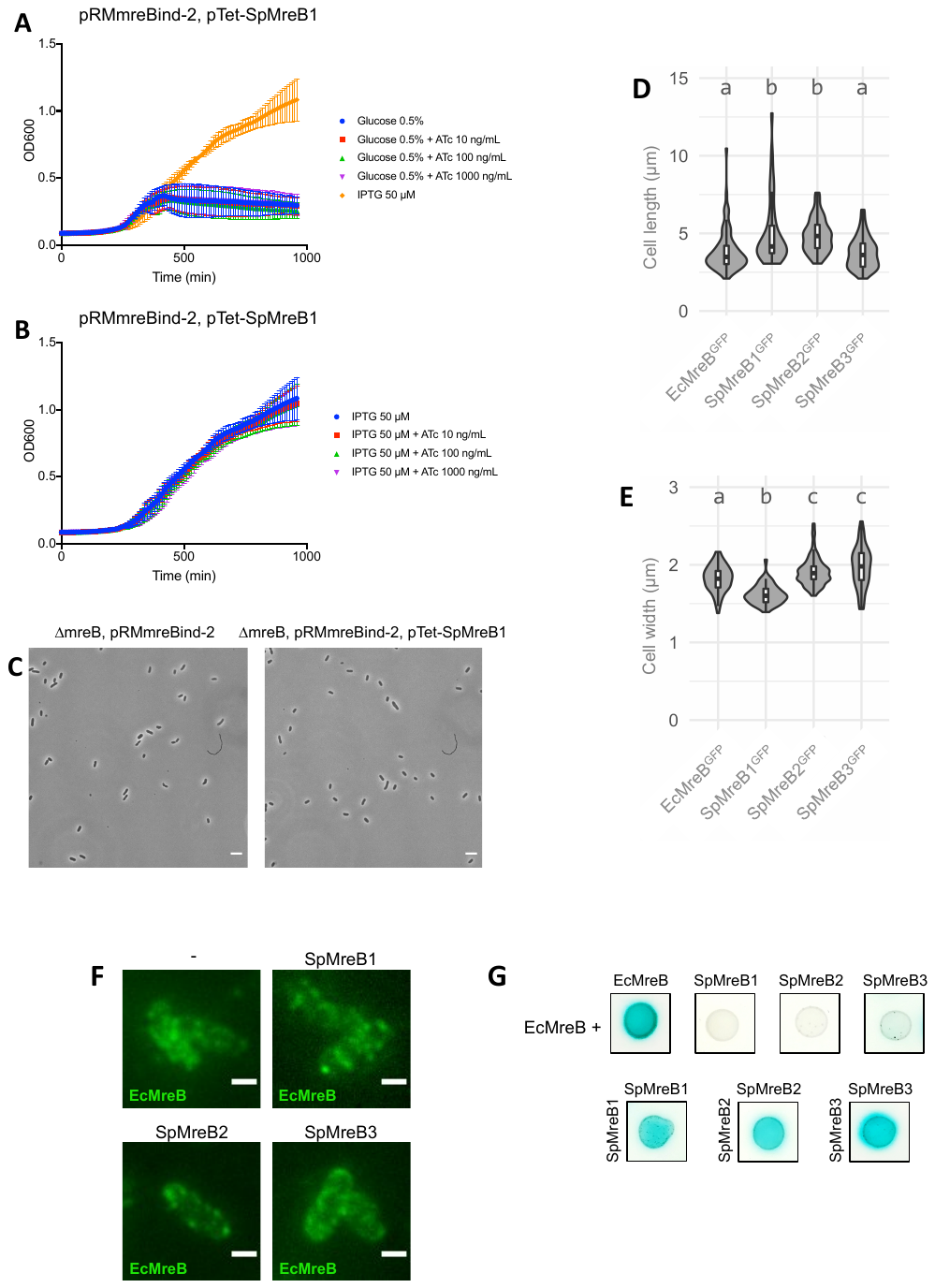
Fig. S2. Growth of *∆mreB* strain complemented with pRMmreBind-2 (IPTG-inducible) and with pTet-SpMreB1 (ATc (anhydrotetracycline)-inducible) in the presence of ATc and glucose (repressed pRM-mreb-ind) or IPTG (induced pRMmreBind-2), showing no rescue of EcMreB function by SpMreB1. (B) Growth of ∆mreB, pRMmreBind-2 cells with 50 µM IPTG and growing concentrations of ATc, showing no toxicity of SpMreB1. (C) Growth of ∆mreB, pRMmreBind-2, pTet-SpMreB1 cells with 50 µM IPTG and 10-1000 ng/mL ATc showing no apparent effect of SpMreB1 expression on cell shape or density. (D) Length and (E) width of cells expressing single untagged SpMreB isoforms. Whisker plots indicate the median and quartiles. Letters above the violin plots indicate the statistical clustering upon ANOVA and post-hoc Tukey-HSD testing. (F) Immunofluorescence images of EcMreB on fixed wild-type cells or cells expressing SpMreB1^GFP^, SpMreB2^GFP^ or SpMreB3^GFP^, indicating that SpMreB production does not alter EcMreB localisation. (G) Two-hybrid interaction in pairwise testing of EcMreB versus SpMreB1, 2 or 3 (top row) and pairwise testing of each isoform against itself (bottom row). A blue color indicates a protein-protein interaction. Each picture represents the strongest signal observed from all pairwise combinations. Scale bars = 2 µm.

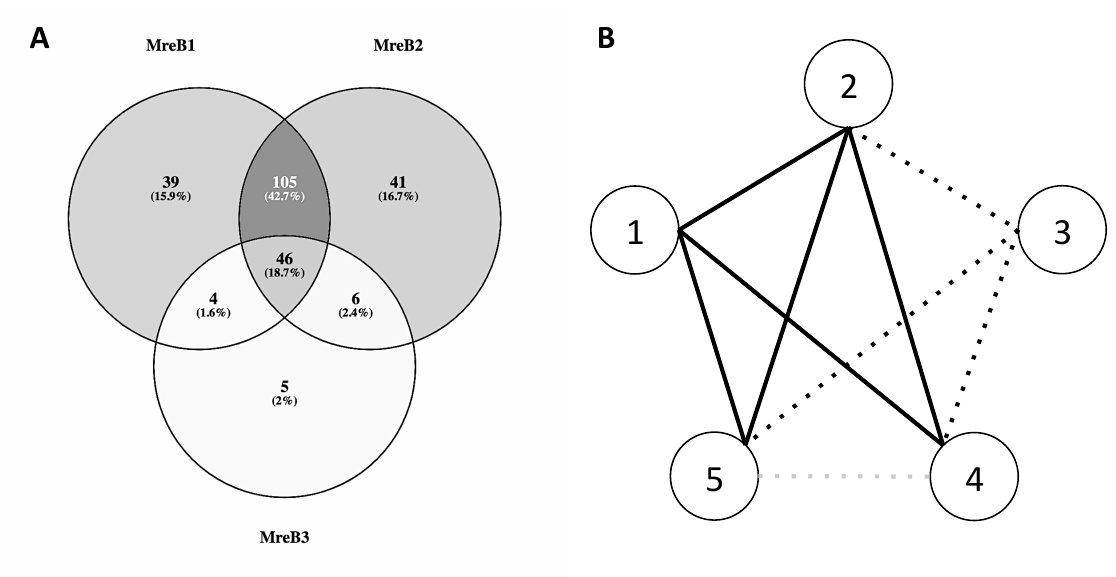

**Fig. S3.** (A) Repartition of the co-immunoprecipitated proteins depending on their interaction with the three bait-constructs (SpMreB1^GFP^, SpMreB1^GFP^, SpMreB1^GFP^ or a combination). (B) Scheme of the inter-isoform interactions predicted from the co-immunoprecipitation data. Numbered circles represent each isoform. Plain lines indicate a strong interaction. Dotted lines indicate a weak interaction (close to significance threshold). The grey dotted line between SpMreB4 and SpMreB5 indicate that this interaction was not be demonstrated experimentally but hypothesized based on the strong homology between SpMreB1 and SpMreB2, and SpMreB4 and SpMreB5, respectively.

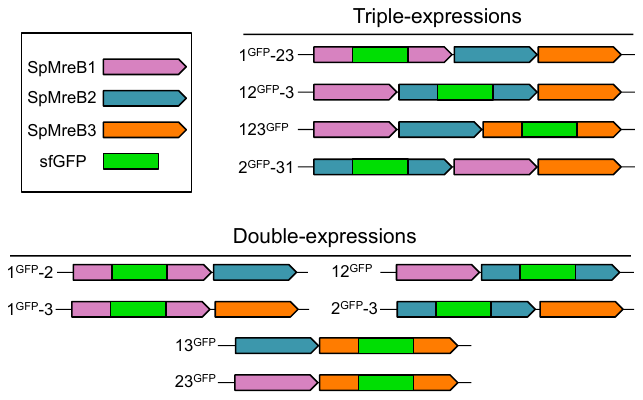

**Supplemental Figure S4 -** Schematics and nomenclature of the constructs used for isoforms coexpression.

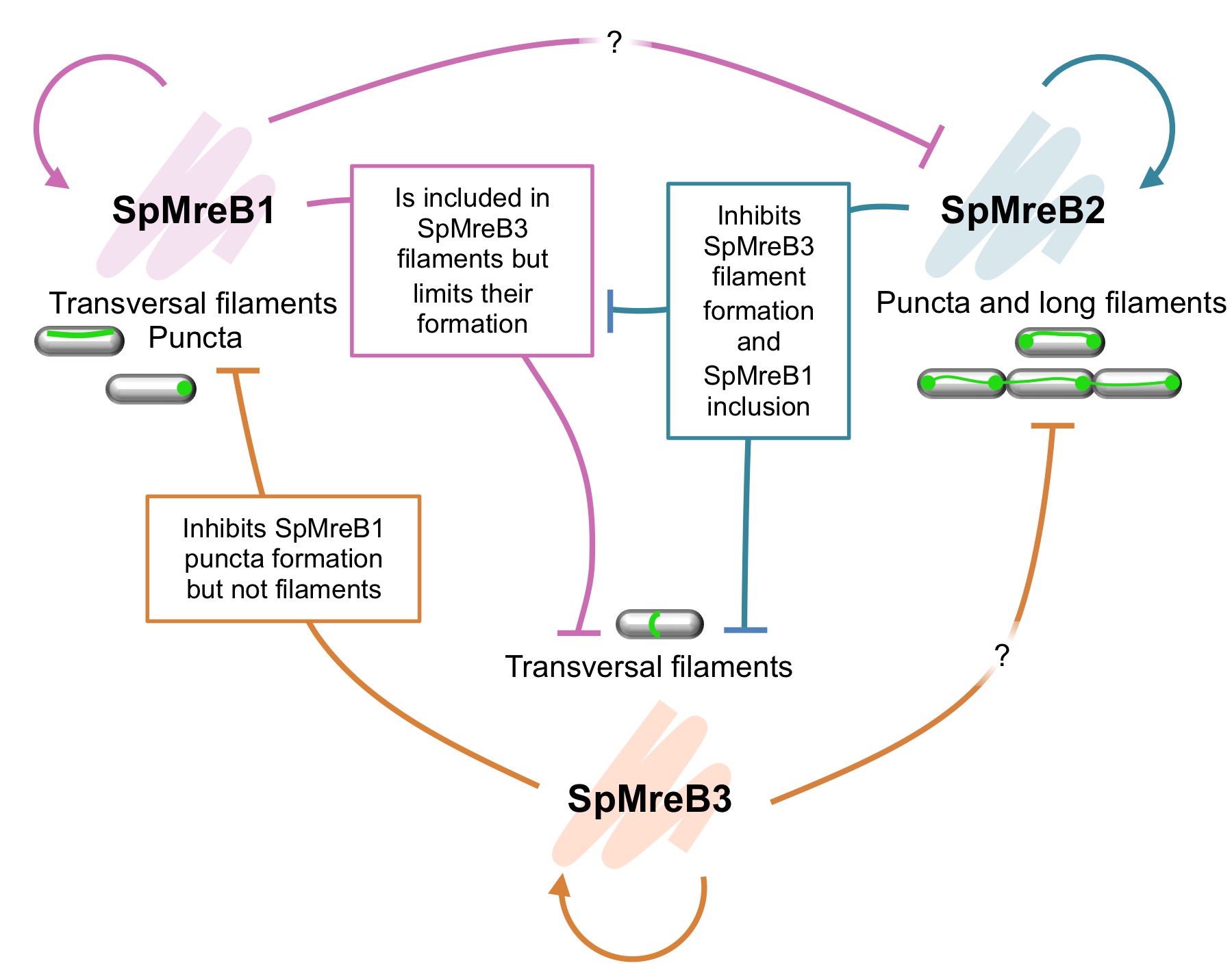

**Figure S5 - Model of SpMreBs interaction network**

Each isoform is capable of forming homopolymers. SpMreB1 forms preferentially straight filaments over the whole length of *E. coli* cells and sometimes forms puncta at one or both poles. SpMreB2 forms large puncta from which protofilaments appear before merging into a thicker single filament that crosses the cells. An overexpression leads to strings of cells that cannot cut the filament to septate properly. SpMreB3 forms transversal filaments. Isoforms coexpressions revealed than SpMreB1 and 3 participate in common structures, while SpMreB2 forms its own. A negative effect of both SpMreB1 and SpMreB2 on SpMreB3 filament formation has been identified. SpMreB2 also represses SpMreB1 filament formation, and SpMreB3 reduces SpMreB1 puncta formation but not filaments.

**Supplementary table 1: Bacterial strains used in this study**

| Species | Strain | Sources |
| --- | --- | --- |
| *Spiroplasma poulsonii* | Ug-1 | (10) |
| *Spiroplasma citri* | GIIX | Gift from UMR1332 “Biologie du Fruit et Pathologie,” INRAE Bordeaux |
| *Spiroplasma melliferum* | KC3 |  |
| *Escherichia coli* | Chemically competent XL10 Gold | Agilent |
| *Escherichia coli* | BL21 | New England Biolabs |
| *Escherichia coli* | BTH101 | Euromedex |
| *Escherichia coli* | MG1655 ∆mreB | Gift from KC Huang (1) |

**Supplementary table 2: Plasmid cloning strategy** (11)

| **Name** | **Code** | **Description, cloning (backbone; primers)** | **Sources** |
| --- | --- | --- | --- |
| pET28(a) | **-** | KanR, IPTG inducible | Novagen |
| pET28-EcMreB^GFP^ | pFM33 | Single expression of *EcMreB* with a sandwich GFP tag (pET28, oFM001, oFM002; synthetic sequence, oFM003, oFM004) | This study |
| pET28-SpMreB1^GFP^ | pXP472 | Single expression of *SpMreB1* with a sandwich GFP tag (pET28; oXP977, oXP978 - synthetic sequence; oXP1043, oXP1041) | This study |
| pET28-SpMreB2^GFP^ | pXP473 | Single expression of *SpMreB2* with a sandwich GFP tag (pET28; oXP977, oXP978 - synthetic sequence; oXP1038, oXP1035) | This study |
| pET28-SpMreB3^GFP^ | pXP474 | Single expression of *SpMreB3* with a sandwich GFP tag (pET28; oXP977, oXP978 - synthetic sequence; oXP1101, oXP982) | This study |
| pET28-SpMreB1^mTur^ | pFM29 | Single expression of *SpMreB1* with a sandwich mTurquoise tag (pET28, oFM001, oFM002; synthetic sequence, oFM005, oFM006) | This study |
| pET28-SpMreB2^mScarlet^ | pFM30 | Single expression of *SpMreB2* with a sandwich mScarlet tag (pET28, oFM007, oFM008; synthetic sequence, oFM009, oFM010) | This study |
| pET28-SpMreB1^G278D/GFP^ | pXP455 | Inactivating point-mutation of pXP472 (pXP472; oXP972, oXP1053, oXP969, oXP1055) | This study |
| pET28-SpMreB2^G279D/GFP^ | pXP470 | Inactivating point-mutation of pXP473 (pXP473; oXP1097, oXP972, oXP973, oXP1098) | This study |
| pET28-SpMreB3^G288D/GFP^ | pXP471 | Inactivating point-mutation of pXP474 (pXP474; oXP1099, oXP972, oXP973, oXP1100) | This study |
| pET28-SpMreB1^GFP^-SpMreB2^mTur^-SpMreB3^mScarlet^ | pXP453 | Co-expression of three isoforms with different tags (pXP426; oXP1045, oXP1046 - pFM29; oXP1047, oXP1048 - pXP427; oXP1049, oXP1050 - pFM30; oXP1051, oXP1052) | This study |
| pET28-1^GFP^-23 | pXP426 | Co-expression plasmid with GFP tag on a single isoform (pET28; oXP977, oXP978 - synthetic construct ; oXP979, oXP980, oXP971, oXP972, oXP981, oXP982) | This study |
| pET28-1^GFP^-2 | pXP445 | Co-expression plasmid with GFP tag on a single isoform (pET28; oXP977, oXP978 - pXP426 ; oXP1034, oXP1035) | This study |
| pET28-1^GFP^-3 | pXP447 | Co-expression plasmid with GFP tag on a single isoform (pET28; oXP977, oXP978 - pXP426 ; oXP1034, oXP1036, oXP1037, oXP982) | This study |
| pET28-12^GFP^-3 | pXP427 | Co-expression plasmid with GFP tag on a single isoform (pET28; oXP977, oXP978 - synthetic construct ; oXP979, oXP983, oXP971, oXP972, oXP984, oXP982) | This study |
| pET28-2^GFP^-3 | pXP448 | Co-expression plasmid with GFP tag on a single isoform (pET28; oXP977, oXP978 - pXP427 ; oXP1038, oXP982) | This study |
| pET28-12^GFP^ | pXP446 | Co-expression plasmid with GFP tag on a single isoform (pET28; oXP977, oXP978 - pXP427 ; oXP1034, oXP1035) | This study |
| pET28-123^GFP^ | pXP428 | Co-expression plasmid with GFP tag on a single isoform (pET28; oXP977, oXP978 - synthetic construct ; oXP979, oXP985, oXP971, oXP972, oXP986, oXP982) | This study |
| pET28-13^GFP^ | pXP451 | Co-expression plasmid with GFP tag on a single isoform (pXP428 ; oXP1038, oXP982) | This study |
| pET28-23^GFP^ | pXP452 | Co-expression plasmid with GFP tag on a single isoform (pXP428 ; oXP1036, oXP1037) | This study |
| pET28-2^GFP^-31 | pXP449 | Co-expression plasmid with GFP tag on a single isoform (pET28; oXP977, oXP978 - pXP427 ; oXP1038, oXP1039, oXP1040, oXP1041) | This study |
| pRMmreBInd-2 | - | Carries a WT copy of *E. coli mreB* (for complementation of the ΔmreB strain). IPTG inducible. | Gift from KC Huang (1) |
| pDSG323 | - | Originally designed for surface-display. KanR, Tet inducible | (11) |
| pDSG323(pTet)-SpMreB1 | pXP419 | Tet-inducible SpMreB1. Marker has been replaced by AmpR. (pDSG323; oFM011, oFM012, oXP965, oXP966 - synthetic construct; oFM013, oFM014 - pUT18; oXP967, oXP968) | This study |
| pDSG323(pTet)-SpMreB2 | pXP420 | Tet-inducible SpMreB1. Marker has been replaced by AmpR. (pDSG323; oFM011, oFM012, oXP965, oXP966 - synthetic construct; oFM015, oFM016 - pUT18; oXP967, oXP968) | This study |
| pDSG323(pTet)-SpMreB3 | pXP421 | Tet-inducible SpMreB1. Marker has been replaced by AmpR. (pDSG323; oFM011, oFM012, oXP965, oXP966 - synthetic construct; oFM017, oFM018 - pUT18; oXP967, oXP968) | This study |
| pUT18 | - | 2-hybrid backbone | Euromedex |
| pKNT25 | - | 2-hybrid backbone | Euromedex |
| pUT18C | - | 2-hybrid backbone | Euromedex |
| pKT25 | - | 2-hybrid backbone | Euromedex |
| pUT18-MreB1_T18 | pXP429 | Construct for 2-hybrid (pUT18; oXP987, oXP988 - pXP419, oXP989, oXP990) | This study |
| pKNT25-MreB1_T25 | pXP430 | Construct for 2-hybrid (pKTN25; oXP991, oXP988 - pXP419, oXP992, oXP993) | This study |
| pUT18C-T18_MreB1 | pXP431 | Construct for 2-hybrid (pUT18C; oXP994, oXP995 - pXP419, oXP996, oXP997) | This study |
| pKT25-T25_MreB1 | pXP432 | Construct for 2-hybrid (pKT25; oXP998, oXP999 - pXP419, oXP1000, oXP1001) | This study |
| pUT18-MreB2_T18 | pXP433 | Construct for 2-hybrid (pUT18; oXP987, oXP988 - pXP420, oXP1002, oXP1003) | This study |
| pKNT25-MreB2_T25 | pXP434 | Construct for 2-hybrid (pKNT25; oXP991, oXP988 - pXP420, oXP1004, oXP1005) | This study |
| pUT18C-T18_MreB2 | pXP435 | Construct for 2-hybrid (pUT18C; oXP994, oXP995 - pXP420, oXP1006, oXP1007) | This study |
| pKT25-T25_MreB2 | pXP436 | Construct for 2-hybrid (pKT25; oXP998, oXP999 - pXP420, oXP1008, oXP1009) | This study |
| pUT18-MreB3_T18 | pXP437 | Construct for 2-hybrid (pUT18; oXP987, oXP988 - pXP421, oXP1010, oXP1011) | This study |
| pKNT25-MreB3_T25 | pXP438 | Construct for 2-hybrid (pKNT25; oXP991, oXP988 - pXP421, oXP1012, oXP1013) | This study |
| pUT18C-T18_MreB3 | pXP439 | Construct for 2-hybrid (pUT18C; oXP994, oXP995 - pXP421, oXP1014, oXP1015) | This study |
| pKT25-T25_MreB3 | pXP440 | Construct for 2-hybrid (pKT25; oXP998, oXP999 - pXP421, oXP1016, oXP1017) | This study |
| pUT18-EcoMreB_T18 | pXP441 | Construct for 2-hybrid (pUT18; oXP987, oXP988 - *E. coli* K12 gDNA; oXP1018, oXP1019) | This study |
| pKNT25-EcoMreB_T25 | pXP442 | Construct for 2-hybrid (pKNT25; oXP991, oXP988 - *E. coli* K12 gDNA; oXP1020, oXP1021) | This study |
| pUT18C-T18_EcoMreB | pXP443 | Construct for 2-hybrid (pUT18C; oXP994, oXP995 - *E. coli* K12 gDNA; oXP1022, oXP1023) | This study |
| pKT25-T25_EcoMreB | pXP444 | Construct for 2-hybrid (pKT25; oXP998, oXP999 - *E. coli* K12 gDNA; oXP1024, oXP1025) | This study |

**Supplementary table 3: Primer table**

| **Name** | **Sequence** |
| --- | --- |
| oFM001 | GGTATATCTCCTTCTTAAAGTTAAACAAAATTATTTC |
| oFM002 | GAGATCCGGCTGCTAACAAAG |
| oFM003 | TAGCAGCCGGATCTCTTACTCTTCGCTGAACAG |
| oFM004 | TTTAAGAAGGAGATATACCATGTTGAAAAAATTTCGTGG |
| oFM005 | TTTAAGAAGGAGATATACCATGGCAATGTTTAATAATAAAAAGC |
| oFM006 | TAGCAGCCGGATCTCTTAATAATCAAGCTCTTTTGTTTTTTG |
| oFM007 | GCTGCTGCCGCTTTTCACCAGCGCGCCAATTTC |
| oFM008 | GCCCGGCGCACCGCTATTATTAATGAAAAACCAGTGCG |
| oFM009 | TAAAAAAGAAATTGGCGCGCTGGTGAAAAGCGGCAGCAGCGTGAGCAAGGGCGAGGCA |
| oFM010 | CACGCACTGGTTTTTCATTAATAATAGCGGTGCGCCGGGCCTTGTACAGCTCGTCCATGCC |
| oFM011 | TAATAATACTAGTAGCGGCCG |
| oFM012 | CTAGTATTTCTCCTCTTTCTCTAGTAG |
| oFM013 | ACTAGAGAAAGAGGAGAAATACTAGATGGCAATGTTTAATAATAAAAAGCC |
| oFM014 | GCAGCGGCCGCTACTAGTATTATTAATAATCAAGCTCTTTTGTTTTTTGATG |
| oFM015 | ACTAGAGAAAGAGGAGAAATACTAGATGGCTAATTATAAATTTGGAAAAGATTATTC |
| oFM016 | GCAGCGGCCGCTACTAGTATTATTAAGCAATTTTGTACTCGTTAGC |
| oFM017 | ACTAGAGAAAGAGGAGAAATACTAGATGGCTATATCTGACGTTCTTAAG |
| oFM018 | GCAGCGGCCGCTACTAGTATTATTAACGGTTTTTTTTATTTTCCTCAATTAATC |
| oXP0965 | GTGGCTTTGTTGAATAAATCGAAC |
| oXP0966 | TCAGAATTGGTTAATTGGTTGTAAC |
| oXP0967 | AACCAATTAACCAATTCTGATTACCAATGCTTAATCAGTGAGG |
| oXP0968 | GATTTATTCAACAAAGCCACGAGCTGCATGTGTCAGAG |
| oXP0969 | ATAAAAGCGGTGCGCCGGGCCATGGTGACGAGCGTAAG |
| oXP0971 | AGCGGCAGCAGCAGC |
| oXP0972 | GCCCGGCGCACCGC |
| oXP0973 | ATAAAAGCGGTGCGCCGGGCATTATTAATGAGAAGCCTGTTCGTG |
| oXP0977 | GATCCGGCTGCTAACAAAG |
| oXP0978 | ACAAAATTATTTCTAGAGGGGAATTGTTATC |
| oXP0979 | CCCTCTAGAAATAATTTTGTTCAGGATTATCCCTTAGTATGGCGATGT |
| oXP0980 | CCTTTGCTGCTGCTGCCGCTTTTCGCCAGGCTACCAATTTC |
| oXP0981 | ATAAAAGCGGTGCGCCGGGCCATGGTGATGAACGTAAAATGAAAG |
| oXP0982 | GCTTTGTTAGCAGCCGGATCTTAGCGATTTTTTTTGTTTTCTTCAATC |
| oXP0983 | CCTTTGCTGCTGCTGCCGCTTTTCACCAGCGCGCC |
| oXP0984 | ATAAAAGCGGTGCGCCGGGCATTATTAATGAAAAACCAGTGCGTG |
| oXP0985 | CCTTTGCTGCTGCTGCCGCTTTTATACAGGCTGCCAATCTG |
| oXP0986 | ATAAAAGCGGTGCGCCGGGCACCAAAGAAAACAAAAAAATGATTATTTTTG |
| oXP0987 | AGCTCGAATTCAGCCGC |
| oXP0988 | GGTCATAGCTGTTTCCTGTGTG |
| oXP0989 | CACAGGAAACAGCTATGACCATGGCAATGTTTAATAATAAAAAGCCTAC |
| oXP0990 | CTGGCGGCTGAATTCGAGCTATAATCAAGCTCTTTTGTTTTTTGATG |
| oXP0991 | AGCTCGAATTCAATGACCATG |
| oXP0992 | CACAGGAAACAGCTATGACCATGGCAATGTTTAATAATAAAAAGCCTAC |
| oXP0993 | ATGGTCATTGAATTCGAGCTATAATCAAGCTCTTTTGTTTTTTGATG |
| oXP0994 | CTAAGTAATATGGTGCACTCTCAG |
| oXP0995 | CTCTAGAGTCGACCTGCAG |
| oXP0996 | TGCAGGTCGACTCTAGAGATGGCAATGTTTAATAATAAAAAGCC |
| oXP0997 | AGTGCACCATATTACTTAGTTAATAATCAAGCTCTTTTGTTTTTTGATG |
| oXP0998 | CTAAGAATTCGGCCGTCG |
| oXP0999 | CTCTAGAGTCGACCCTGC |
| oXP1000 | CTGCAGGGTCGACTCTAGAGATGGCAATGTTTAATAATAAAAAGCC |
| oXP1001 | ACGACGGCCGAATTCTTAGTTAATAATCAAGCTCTTTTGTTTTTTGATG |
| oXP1002 | AGGAAACAGCTATGACCATGGCTAATTATAAATTTGGAAAAGATTATTC |
| oXP1003 | GAATTCGAGCTCGGTACCCGAGCAATTTTGTACTCGTTAGCC |
| oXP1004 | AGGAAACAGCTATGACCATGGCTAATTATAAATTTGGAAAAGATTATTC |
| oXP1005 | ATGGTCATTGAATTCGAGCTAGCAATTTTGTACTCGTTAGCC |
| oXP1006 | GCAGGTCGACTCTAGAGATGGCTAATTATAAATTTGGAAAAGATTATTC |
| oXP1007 | GAGTGCACCATATTACTTAGTTAAGCAATTTTGTACTCGTTAGC |
| oXP1008 | CAGGGTCGACTCTAGAGATGGCTAATTATAAATTTGGAAAAGATTATTC |
| oXP1009 | AACGACGGCCGAATTCTTAGTTAAGCAATTTTGTACTCGTTAGC |
| oXP1010 | CACAGGAAACAGCTATGACCATGGCTATATCTGACGTTCTTAAG |
| oXP1011 | GAATTCGAGCTCGGTACCCGACGGTTTTTTTTATTTTCCTCAATTAATC |
| oXP1012 | CACAGGAAACAGCTATGACCATGGCTATATCTGACGTTCTTAAG |
| oXP1013 | ATGGTCATTGAATTCGAGCTACGGTTTTTTTTATTTTCCTCAATTAATC |
| oXP1014 | ACTGCAGGTCGACTCTAGAGATGGCTATATCTGACGTTCTTAAG |
| oXP1015 | TGCACCATATTACTTAGTTAACGGTTTTTTTTATTTTCCTCAATTAATC |
| oXP1016 | CTGCAGGGTCGACTCTAGAGATGGCTATATCTGACGTTCTTAAG |
| oXP1017 | GACGGCCGAATTCTTAGTTAACGGTTTTTTTTATTTTCCTCAATTAATC |
| oXP1018 | CACAGGAAACAGCTATGACCATGTTGAAAAAATTTCGTGGCATG |
| oXP1019 | GAATTCGAGCTCGGTACCCGCTCTTCGCTGAACAGGTCG |
| oXP1020 | CACAGGAAACAGCTATGACCATGTTGAAAAAATTTCGTGGCATG |
| oXP1021 | ATGGTCATTGAATTCGAGCTCTCTTCGCTGAACAGGTCG |
| oXP1022 | ACTGCAGGTCGACTCTAGAGATGTTGAAAAAATTTCGTGGCATG |
| oXP1023 | GAGTGCACCATATTACTTAGTTACTCTTCGCTGAACAGGT |
| oXP1024 | CTGCAGGGTCGACTCTAGAGATGTTGAAAAAATTTCGTGGCATG |
| oXP1025 | AACGACGGCCGAATTCTTAGTTACTCTTCGCTGAACAGGTC |
| oXP1034 | CCCTCTAGAAATAATTTTGTTCAGGATTATCCCTTAGTATGGCGATGTT |
| oXP1035 | CTTTGTTAGCAGCCGGATCTTACGCAATTTTATATTCATTCGCTTCCAG |
| oXP1036 | TTAATAATCCAGTTCTTTGGTTTTCTGATGTTC |
| oXP1037 | CCAAAGAACTGGATTATTAATCTAGAGCATGCGTCGAC |
| oXP1038 | CCTCTAGAAATAATTTTGTTCAGGATTATCCCTTAGTATGGCGAATTAT |
| oXP1039 | ATCTAGACCATGGGAATTCTTAGCGATTTTTTTTGTTTTCTTCAATCAG |
| oXP1040 | GAATTCCCATGGTCTAGATCAGGATTATCCCTTAGTATGGC |
| oXP1041 | GCTTTGTTAGCAGCCGGATCTTAATAATCCAGTTCTTTGGTTTTCTG |
| oXP1043 | CTAGAAATAATTTTGTTCAGGATTATCCC |
| oXP1045 | CGGGCACCAAAGAAAACAAAAAAATGATTATTTTTG |
| oXP1046 | TTTCACCAGCGCGCC |
| oXP1047 | AAATTGGCGCGCTGGTGAAAGGTGGTGGGGGCGG |
| oXP1048 | TGGTTTTTCATTAATAATAGCAGTACGACGAGCC |
| oXP1049 | GCTCGTCGTACTGCTATTATTAATGAAAAACCAGTGCGTG |
| oXP1050 | GACGCTGGACCCGGATTTATACAGGCTGCCAATCTG |
| oXP1051 | AGATTGGCAGCCTGTATAAATCCGGGTCCAGCGTC |
| oXP1052 | TTTGTTTTCTTTGGTGCCCGGCGCACCGCTCTTGTAAAGTTCATCCATACCACC |
| oXP1053 | GACATCTTCCGCAACGGT |
| oXP1055 | GTTGCGGAAGATGTCATCAGCCAATTCTGGAG |
| oXP1097 | GATGTTATCCGCAATGGAATCAC |
| oXP1098 | ATTGCGGATAACATCATCGGCAAGCTCAGGGG |
| oXP1099 | GATGCTGTTGCTAATGGAATTTTAGTTAC |
| oXP1100 | ATTAGCAACAGCATCATCAGCAAGCTCTGCTGG |
| oXP1101 | CTAGAAATAATTTTGTTCAGGATTATCCCTTAGTATGGCG |

Dataset S1 (separate file). LC-MS/MS count table from SpMreBs co-immunoprecipitation.

Movie S1 (separate file). Representative timelapse of *S. poulsonii* motility.

Movie S2 (separate file). Timelapse of SpMreB2^GFP^ protofilament merging.

Movie S3 (separate file). Timelapse of SpMreB2^GFP^ protofilament detaching from puncta upon cell elongation.
